# High red/far-red ratio promotes root colonization of *Serratia plymuthica* A21-4 in tomato by root exudates-stimulated chemotaxis and biofilm formation

**DOI:** 10.1101/2023.07.06.547930

**Authors:** Zhixin Guo, Yanping Qin, Jingli Lv, Xiaojie Wang, Ting Ye, Xiaoxing Dong, Nanshan Du, Tao Zhang, Fengzhi Piao, Han Dong, Shunshan Shen

**Affiliations:** College of Horticulture, Henan Agricultural University, Zhengzhou 450002, P. R. China; College of Plant Protection, Henan Agricultural University, Zhengzhou 450002, China

**Keywords:** plant growth-promoting rhizobacteria, light quality, metabolomics, root exudate compontents, colonization ability

## Abstract

Effective colonization on plant roots is a prerequisite for plant growth promoting rhizobacterias (PGPR) to exert beneficial activities. Light is essential for plant growth, development and stress response. However, how light modulates root colonization of PGPR remains unclear. Here, we found that high red/far red(R/FR) light promoted and low R/FR light inhibited the colonization and growth enhancement of *Serratia plymuthica* A21-4 on tomato. Non-targeted metabolomic analysis of root exudates collected from different R/FR ratio treated tomato seedlings with A21-4 inoculation by ultra performance liquid chromatography-tandem mass spectrometry showed that 64 primary metabolites including amino acids, sugars and organic acids in high R/FR light-grown plants significantly increased compared with those determined for low R/FR light-grown plants. Among them, 7 amino acids, 1 organic acid and 1 sugar obviously induced the chemotaxis and biofilm formation of A21-4 compared to the control. Furthermore, exogenous addition of five artificial root exudate compontents (leucine, methionine, glutamine, 6-aminocaproic acid and melezitose) regained and further increased the colonization and growth promoting ability of A21-4 in tomato under low R/FR light and high R/FR light, respectively, indicating their involvement in high R/FR light-regulated the interaction of tomato root and A21-4. Taken together, our results, for the first time, clearly demonstrate that high R/FR light-induced root exudates play a key role in chemotaxis, biofilm formation and root colonization of A21-4. This study provides new insights into the interactions of plant-PGPR under different light conditions and can help promote the combined application of light supplementation and PGPR to facilitate crop growth and health in green agricultural production.

## 1. Introduction

Global food production is estimated to increase by 70% by 2050 to meet the increasingly affluent world population that is predicted to exceed 9. 5 billion(Haskett et al, 2021). However, agricultural production, as a major contributor to global food supply, is highly dependent on crops which are seriously threatened by changing climatic conditions, abiotic and biotic stresses and poor soil quality(Zhang et al, 2022). Ensuring food security is achieved generally through the overuse of pesticides and fertilizers; However, their application often leads to pollution of the ecosystem(Mburu et al, 2022). Therefore, some novel and environmentally friendly aspects of agricultural practices are urgently required to solve the food problems of the future and protect ecosystems. As a simple, sustainable and environmental-friendly method, application of plant growth-promoting rhizobacteria (PGPR) can enhance plant resistance to various biotic and abiotic stresses and reduce pollution from fertilizers and pesticides(Majeed et al, 2018).

PGPR is a collective term for a group of rhizosphere-inhabiting beneficial bacteria capable of facilitating plant growth and health(Shameer and Prasad, 2018). PGPR include many diverse bacterial genera such as *Alcaligenes*, *Arthrobacter*, *Bacillus*, *Clostridium*, *Derxia*, *Enterobacter*, *Flavobacterium*, *Gluconacetobacter*, *Herbaspirillum*, *Pseudomonas*, *Rhodococcus*, *Serratia* and *Zoogloea*(Cook et al, 1995). The mechanisms by which PGPR exert their beneficial effects consist of dissolving potassium; solubilizing phosphorus; fixing nitrogen; producing bioactive volatile organic compounds, siderophores, gibberellic acid, indoleacetic acid, and antimicrobial substances; modifying the composition and functioning of rhizosphere microbial community; and triggering induced systemic resistance against pathogens(Chandran et al, 2021; Vejan et al, 2016). Root colonization is critical for PGPR to exert their beneficial effects on plants, however, the insufficient colonization of PGPR on the roots of crops is a widespread problem severely limiting the application of PGPR in the field(Shameer and Prasad, 2018). Hence, it is pivotal to understand the processes of the root colonization of PGPR.

Chemotaxis and biofilm formation are vital for root colonization and for exerting the beneficial functions of PGPR(dos Santos et al, 2020). Chemotaxis initiated by ligand sensing at methyl-accepting chemotaxis proteins (MCPs) covers bacteria being induced and recruited by specific root exudate components to gather around plant roots. Thus, chemotaxis is a prerequisite for PGPR to colonize plant roots(Xiong et al, 2020; Yaryura et al, 2008). Biofilm formation is also an important factor affecting the colonization ability of PGPR. Biofilms enhance the resistance of bacteria to environmental stress and improve the interaction between bacteria and various abiotic or biotic surfaces(Liu et al, 2020), thereby resulting in the efficient colonization of PGPR on different surfaces. Moreover, stable biofilm on the plant root surfaces is an important indication of successful PGPR colonization.

Plant root exudates and its compositions play an important role in inducing the chemotactic response and biofilm formation of PGPR. For instance, cucumber root-released D-galactose functioned as a strong chemoattractant and significantly enhanced the biofilm formation of *Bacillus velezensis* SQR9(Liu et al, 2020). Amino acids, organic acids and sugars are the majority components of primary metabolites in root exudates(Gao et al, 2023). Plants secrete up to 10–40% of their photosynthetically fixed carbon into the rhizosphere as root exudates(Korenblum et al, 2020; Vives-Peris et al, 2020). The quality and quantity of root exudates may vary among plant species, by plant age, and by environmental factors such as light(Gundel et al, 2014).

Light is essential for plants both a source of energy for photosynthesis and a vital environmental signal that controls plant growth, development and response to stress via specific sensory photoreceptors(Petroutsos et al, 2016). Light quality usually involves the R/FR light ratio detected by the phytochromes(Lim et al, 2018). Plants, either in nature or in agriculture, are often exposed to low R/FR light which can occur because of high density and the transition from warm to cool seasons(Wang et al, 2018). However, to date, how light regulates root colonization of plants by PGPR remains elusive. Interestingly, some reports have shown that light affects plant root exudates, suggesting that light may also influence the root colonization of PGPR. For example, low light altered root exudation and reduced putative beneficial microorganisms in seagrass roots(Martin et al, 2018). Enhanced UV-B exposure reduced allocation to the below-ground biomass and root exudation of the mire plants and induced a decrease in the contents of malic acid and tartaric acid and an increase in the contents of succinic acid and oxalic acid in rice root exudates, respectively(He et al, 2016; Rinnan et al, 2006).

*Serratia plymuthica* A21-4, as a promising biocontrol agent, was isolated from the root of onion (*Allium cepa* L.)(Shen et al, 2002)and the strain showed high efficacy on growth promoting activity(Shen et al, 2005)and the control of Phytophthora blight of pepper(Shen et al, 2007). Nevertheless, the role of light in regulating the root colonization of the strain A21-4 in crops remains unclear. To verify the hypothesis that R/FR ratio is pivotal for colonization of the strain A21-4, we first compared the growth promoting effect and root colonization capability of A21-4 on the tomato plants under different R/FR ratio conditions, then explored root exudates changes of tomato seedlings with or without A21-4 inoculation under different R/FR light ratios through untargeted metabolic profiling based on UPLC-MS/MS, and finaly evaluated the effects of specific differential root exudate components on the chemotaxis, biofilm formation and root colonization of A21-4. Based on our results, we confirmed that R/FR light ratio-induced root exudates play a key role in root colonization of A21-4 on tomato seedlings.

## 2. Materials and methods

### 2.1 Plant and bacterial strain materials and growth conditions

Tomato (*Solanum lycopersicum* cv Moneymaker) was used as plant material. Tomato plants were grown in sterilized vermiculite and turf (10:1) and irrigated every four days with half Hoagland’s nutrition solution. Seedlings were planted in chambers until the 3-leaf stage with the following environmental conditions:humidity of 70 %, temperature of 25/20 °C (day/night), 12-h photoperiod, and white light PPFD of 280 µmol m^-2^s^-1^. *Serratia plymuthica* A21-4 strain labeled with rifampicin resistance was used as model strain. A21-4 was stored at −70 °C in TSB containing 20 % glycerol for reserve and grown at 28 °C in TSA medium (TSB 30 g·L^-1^ and agar powder 18 g·L^-1^) for 2-3 days and prepared into 10^8^ cfu·mL^-1^ bacterial suspension with distilled water for later use.

### 2.2 Experimental design and treatments

Three different experiments were carried out. Experiment 1 was conducted to determine the impacts of light quality on the growth enhancement and root colonization of A21-4 in tomato seedlings. Tomato seedlings with three leaves were pretreated low (L, 1:2. 5) or high (H, 2. 5:1) red (R, 660 nm) /far-red (FR, 730 nm) ratios with 12-h photoperiod and PPFD of 140 μmol m^-2^s^-1^ for 3 days and then plants were inoculated A21-4 using root irrigation(100 mL bacterial suspension at a concentration of 10^8^ cfu·mL^-1^ for each tomato plant, and corresponding volume of distilled water was used as control. After that, seedlings with or without A21-4 inoculation were continuously exposed to the same high or low R /FR ratio for 10 days.

Experiment 2 was performed to collect root exudates of different treatments. Tomato plants were grown on half Hoagland’s nutrient solution under white light until the 3-leaf stage. After that, tomato seedlings were planted to 700 ml pots filled with a mixture of vermiculite and turf (10:1) under low or high R/FR ratio conditions. After 3 day, A21-4 was inoculated at a concentration of 10^8^ cfu·mL^-1^ for each tomato plant and corresponding volume of distilled water was used as control. After 1 day, tomato plants were removed carefully from cultivation medium, and the roots were carefully washed with distilled water to remove adhesive substrates. Finally, the seedlings were placed in a hydroponic box filled with 4 L of half Hoagland’s nutrition solution to collect the root exudates.

Experiment 3 was carried out to analyse the impacts of different root exudate components on low or high R/FR ratio-mediated the growth enhancement and root colonization of A21-4 in tomato seedlings. Both light quality and A21-4 treatments were same as experiment 1, and the difference is that 100 mL different root exudate components (100 mmol·L^-1^) were watered in each tomato seedling, 12 hours prior to inoculation with A21-4.

### 2.3 Determination of growth indicators

Plant height and fresh were detected. Concretely, plant height was measured from the base to shoot apex; fresh weight was determined by an electronic balance.

### 2.4 Determination of the colonization ability of A21-4 in tomato roots

Rifampicin labeling method was detected as described previously(Shen et al, 2007). When measuring the colonization density of A21-4 in tomato roots, the substrate culture bowl was gently removed, then the root substrate was washed in running water, and finally, the water was dried with filter paper. The fresh weight of tomato plants was weighed and recorded. The root samples were ground and added with distilled water, diluted to a certain concentration, coated on 1/10 TSA medium plate containing 50 μg·mL^-1^ rifampicin, and incubated in an incubator at 28 °C for 3-5 days. Colony formation of A21-4 was detected, and the root colonization density was calculated. Colonization of growth-promoting bacteria(cfu·g^-1^)= (Number of colonies in the dish × dilution ratio)/(Amount of bacteria adding liquid in the dish (mL) × root weight (g)).

### 2.5 Root exudates collection, sample preparation and extraction and metabolomic analysis by UPLC–MS/MS

Tomato plants with or without A21-4 inoculation were grown on half Hoagland’s nutrient solution under low or high R/FR ratio conditions for 1 day, and then the nutrient solutions were were collected, lyophilized and stored at −80 °C. Each treatment contained five replicates and each replicate included four plants. For sample preparation and extraction, the samples were thawed from the refrigerator at −80 °C and mixed with vortex for 10 s. 23 mL of the sample was taken after mixing, placed in 50mL of the centrifuge tube, then immersed in liquid nitrogen as a whole and put into the lyophilizer until the samples were completely frozen. After that the samples were dissolved with 0. 5 ml 70% methanol internal standard extract, scrolled for 3 min and then centrifuged (12000 r/min, 4 °C) for 10 min. The supernatant was filtered with a microporous filter membrane (0. 22 μm) and stored in a sample flask for ultra performance liquid chromatography-tandem mass spectrometry (UPLC-MS/MS) test. The primary metabolome of tomato root exudates was performed using the non-targeted metabolome method at Metware Biotechnology Co., Ltd. (Wuhan, Hubei, China) as recently reported(Lin et al, 2023).

### 2.6 Chemotaxis assay

The qualitative chemotactic response of A21-4 towards the selected tomato root exudate components was performed using the drop assay method (Feng et al, 2018)with slight modifcations. Briefly, A21-4 was cultured in TSB medium at 28 °C with shaking at 180 rpm until the cell density (OD600) reached about a value of 1. 0. Then the 45 mL of cell suspensions extracted by centrifugation (25 °C, 10 min, 5000 rpm) were resuspended in the 12 mL of chemotaxis buffer (K2HPO4 4. 4 g·L^-1^, KH2PO4 4. 2 g·L^-1^, EDTA-Na2 7. 45 mg·L^-1^, pH 7. 0) and were added to 4 mL of 1% hydroxypropyl methy cellulose. 4 mL of the mixed suspension was added into a 60-mm-diameter dish, and artificial root exudate component (10 μL for a drop, 0. 1 M) was dropped into the center of the culture dish, with 10 μL chemotaxis buffer as control treatment. Then the culture dish was placed at room temperature and the formation of chemotactic cycle was observed.

The same volume of chemotactic buffer was used to suspend the thalli and 100 μL was absorbed with a 200 μL pipette for further use. A 1 mL syringe was used as the capillary for chemotaxis quantitative test, and 100 μL of 100 mmol **·** L^-1^ the purchased commercial root exudate components were absorbed, respectively, with chemotactic buffer added as control, and repeated three times. Then, the needle of the syringe was inserted into the thin mouth of the above gun head to make the two fully contact, and the experiment was carried out in the ultra-clean work table. The solution in the needle was coated with plate dilution method for bacteria count after standing at room temperature for two hours, and cultured at 28 °C for 2-3 days. After that, the plates were observed, photographed and recorded. The relative chemotaxis index (RCI) was used to describe the chemotactic ability of the strain to root exudates, and was the ratio of the average CFUs treated to the control CFUs. Generally, if RCI ≥ 2, it was considered that the chemotaxis of the treatment was significantly different from that of the control CFUS(Ling et al, 2011).

### 2.7 Biofilm formation assay

Crystal violet staining test was used for quantitative analysis of A21-4 biofilm as described previously(Wang et al, 2019)with slight modifcations. 1 mL TSB medium was added to the 48-well cell culture plate, and 10 μL of the above bacterial solution and different root exudates were added to the 48-well plate which then was placed at 28 °C for 5 days. The medium was sucked out and cleaned 3 times with 1 mL chemotactic buffer solution per well. The chemotactic buffer solution was sucked out, 1 mL methanol was added to each well and fixed for 15 min. After air drying, 1 mL 1% crystal violet was put into each well and dyed at room temperature for 5 min. 1 mL of 33% glacial acetic acid was put into each well and cultured at room temperature for 30 min to dissolve crystal violet. OD value was detected with an enzyme label at 590 nm. The assay was repeated three times, with chemotactic buffer as control.

### 2.8 Statistical analysis

The experiments were conducted in a completely randomized block design with three replicates(each containing eight seedlings), except for experiment 2 with five replicates. Significance was evaluated with one-way analysis of variance (ANOVA). Significant differences denoted by different letters between means were analyzed by Turkey’s test (*P*<0. 05). Metabolite data were log2-transformed for statistical analysis to improve normality and were normalized. Metabolites from 20 samples were used for principal component analysis (PCA), and orthogonal partial least squares discriminant analysis (OPLS-DA) to analyze the variation of root exudates. The fold change and p values were set to 2. 0 and 0. 05, respectively.

## 3. Results

### 3.1 High red/far-red ratio induces the root colonization and growth promoting capability of A21-4 in tomato seedlings

To explore how light affects the root colonization and growth promoting effect of A21-4 in tomato seedlings, we first compared the growth promoting effect of A21-4 on the wild-type tomato plants under different red/far-red ratios conditions. As shown by the phenotype, A21-4 inoculation induced a significant increase in plant fresh weight and height of high R/FR light-treated plants. However, A21-4 inoculation failed to increase the growth of low R/FR light-treated seedlings (Fig. 1A-C). Consistent with these changes, high R/FR light-treated seedlings maintained high populations of A21-4 in the roots, while populations of A21-4 markedly reduced in low R/FR light-treated seedlings (Fig. 1D). Our results suggest that R/FR ratio plays key roles in the root colonization and growth promoting capability of A21-4 in tomato seedlings.

**Figure. 1.**
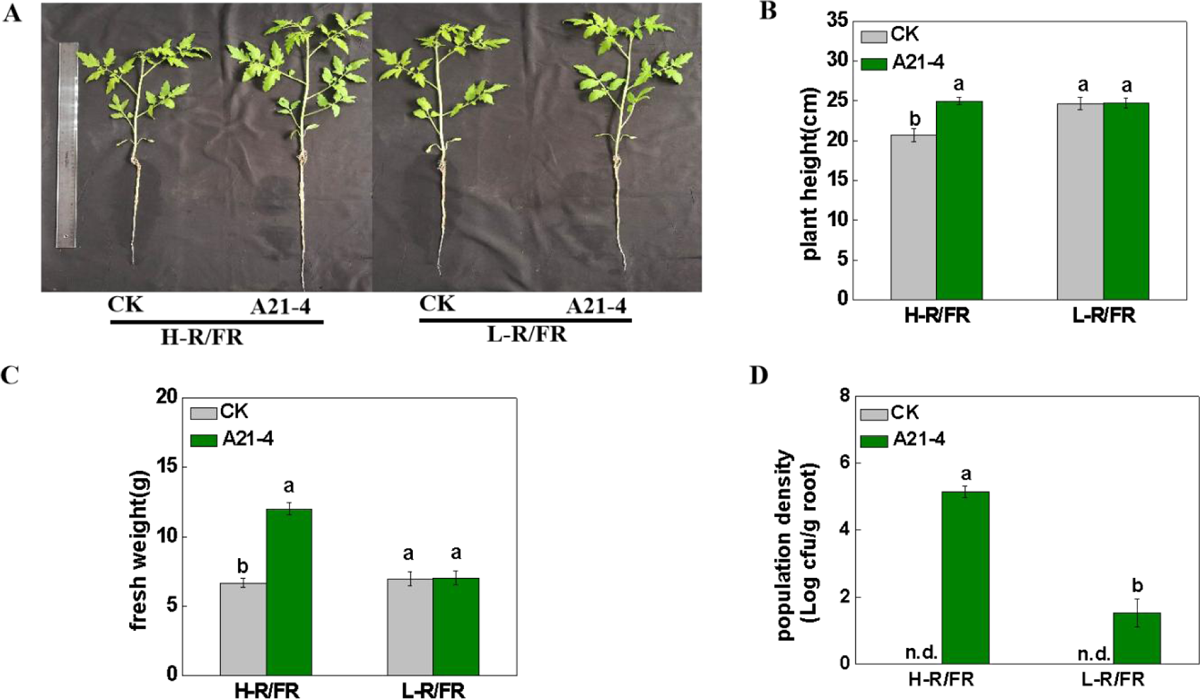
Effect of red/far-red ratios on root colonization and plant growth promotion by *S. plymuthica* A21-4 in tomato seedlings. A-D. Plant phenotype, plant height, fresh weight and root population density of *S. plymuthica* A21-4 of the tomato plants grown under H-R/FR (2. 5:1) or L-R/FR (1:2. 5) conditions at 10 dpi with (A21-4) or without (CK) inoculation. Values are the means ±SD (n=3). The different letters indicate significant difference in the means (*P*< 0. 05, Tukey’s test).

### 3.2 Non-targeted metabolomic analysis of root exudates collected from different R/FR light ratio treated tomato seedlings with or without A21-4 inoculation

Given the predominant role of root exudates in the root colonization of PGPR, non-targeted metabolic profiling based on HPLC-MS/MS was used to study root exudation profile of tomato seedlings with or without A21-4 inoculation under different R/FR light ratios. According to the intra-group repeatability and inter-group discrimination in the PCA score plot, the H-R/FR, H-R/FR+A21-4 and L-R/FR+A21-4 treatment groups had a clear separation, implying that the variation of root exudates induced by A21-4 inoculation and R/FR light ratios. However, the sample points of L-R/FR and L-R/FR+A21-4 treatment groups overlapped partially, suggesting that the content and composition of the root exudates in some samples were similar and A21-4 inoculation led to little change in the variation of the root exudates under low R/FR ratio(Fig. 2A). To detect the differences between the root exudates of tomato seedlings with A21-4 inoculation under different R/FR light ratios to the greatest extent, OPLS-DA was carried out (Fig. 2B).

**Figure. 2.**
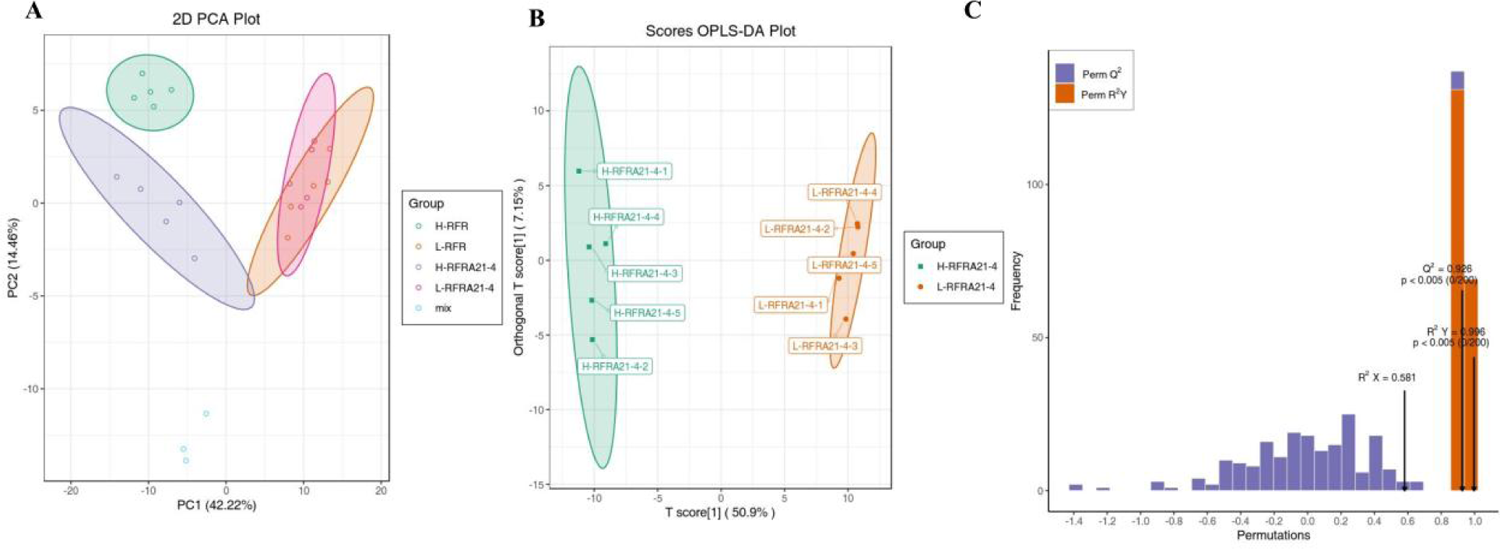
Principal component analysis (PCA) and orthogonal partial least squares discriminant analysis (OPLS-DA) of tomato root exudates primary metabolome. A. PCA score scatter plot depicting root exudate profiles of the tomato plants with or without A21-4 inoculation under H-R/FR (2. 5:1) or L-R/FR (1:2. 5) conditions. B. Orthogonal Projections to Latent Structures Discriminant Analysis (OPLS-DA) score plot on root exudate profiles of the tomato plants with A21-4 inoculation under H-R/FR (2. 5:1) or L-R/FR (1:2. 5) conditions. Each point represents a biological replicate. C. R2 and Q2 represent goodness of fit and prediction respectively, and p-value shows the significance level of the model (x axis=predictive components, y axis=orthogonal component).

Importantly, cross-validation of the OPLS-DA model was excellent with a cumulative R^2^ X of 0. 591, R^2^ Y of 0. 996 and Q^2^ of 0. 926 (Fig. 2C). In the OPLS-DA score chart, the sample points under the same treatment were clearly clustered together, and the sample points between different treatments were obviously separated (Fig. 2B), implying that the model could reflect very well the differences between different treatments and R/FR light ratios exerted an important influence on the quantity and composition of root exudates after A21-4 inoculation.

Furthermore, the Venn diagram showed that 64 common differential metabolites were more abundant in root exudates of tomato colonized by A21-4 under high R/FR light than those in exudates of low R/FR light-treated plant roots after A21-4 inoculation (Fig. 3A). Concretely, to clearly show the changes of root exudates after A21-4 inoculation under different R/FR light ratios, three heat maps were then plotted to visualise the changes in differential metabolites containing organic acids, amino acids and others including sugars (Fig. 3B-D). Taken together, there was a clear change of the metabolite profile of root exudates after A21-4 inoculation under different R/FR light ratios, suggesting that the identified metabolites may play significant roles in the high R/FR light-induced root colonization and growth promoting effect of A21-4 in tomato seedlings.

**Figure. 3.**
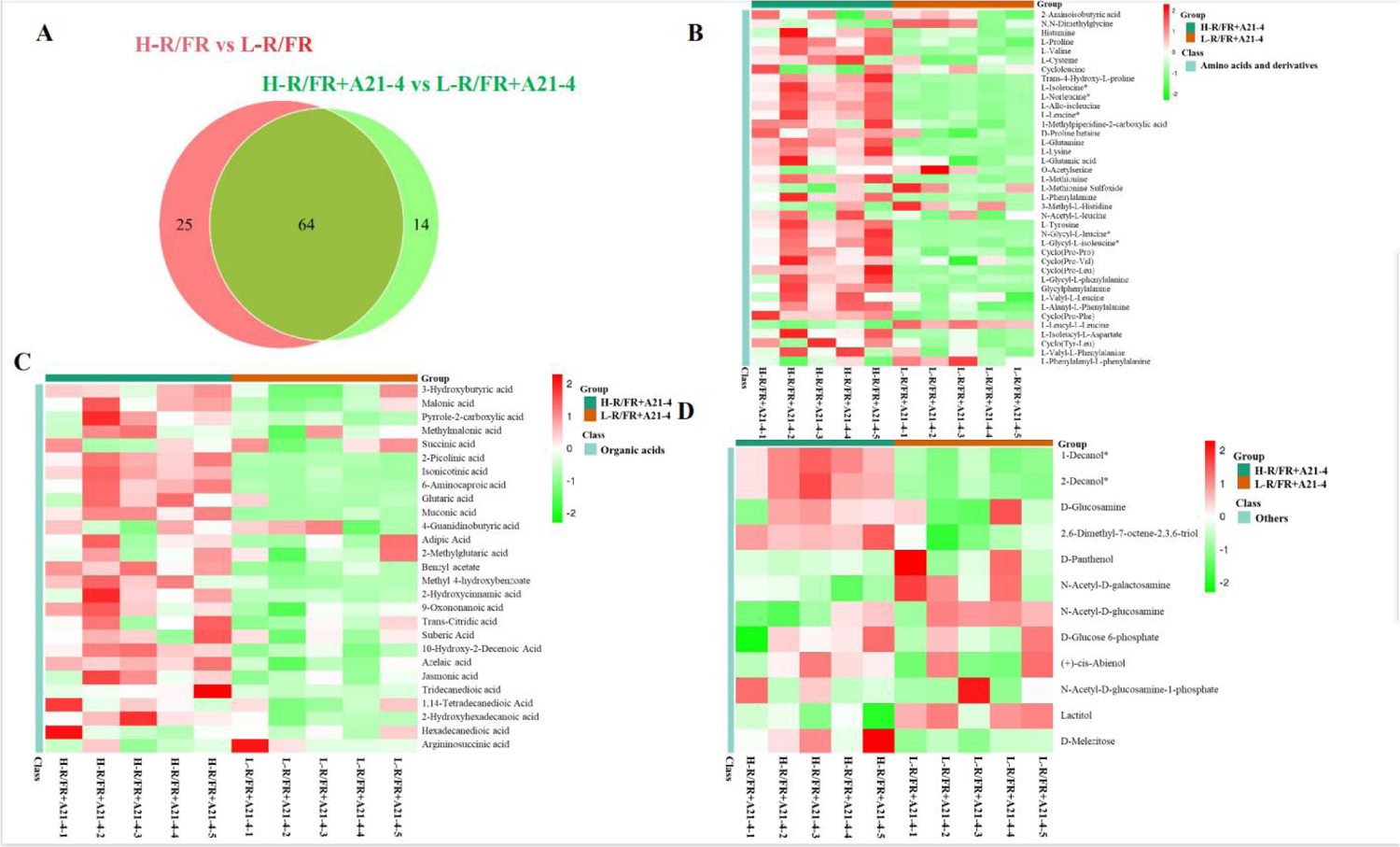
Venn diagram and heat maps of tomato root exudate components effected by A21-4 inoculation and red/far red light ratio A. Venn diagram analysis of tomato root exudate components effected by A21-4 inoculation and red/far red light ratio. B-D. Heatmap analysis of changes of amino acids, organic acids, and others including sugars in root exudate content of A21-4-inoculated plants growing on high and low red/far red light ratio.

### 3.3 Chemotaxis of A21-4 towards selected tomato root exudate components

Since amino acids, organic acids and sugars play key roles in inducing chemotaxis of PGPR, among the identified differential metabolites in root exudates, we selected 10 significant up-regulated root exudate components to evaluate the chemotaxis response of A21-4, considering the availability of different commercially root exudate components and repeatability between different samples. The selected root exudate components contained 7 amino acids(L-Methionine, L-Leucine, L-Glutamine, L-Norleucine L-Isoleucine, L-Lysine and L-Valine), 2 organic acids(6-Aminocaproic acid and 2-Picolinic acid) and 1 sugar(D-Melezitose). The chemotactic response of A21-4 towards the selected root exudate components was determined qualitatively (drop plate assay) and quantitatively (capillary assay), respectively. The results showed that except for 2-Picolinic acid, the rest of 9 root exudate components efficiently induced the chemoattractant activity of A21-4, resulting in the obvious and large bacterial ring in the center of the plate compared to the control (Fig. 4). Interestingly, the chemotactic response of A21-4 towards the selected root exudate components was further confirmed quantitatively by capillary assay. L-Glutamine exhibited the most significant attraction for A21-4 at a 100 mM concentration, followed by 6-Aminocaproic acid, D-Melezitose, L-Leucine, L-Methionine, L-Norleucine, L-Isoleucine, L-Lysine, and L-Valine. However, the effect of 2-Picolinic acid was also not different from the control(Fig. 5). Together, these results suggested that H-R/FR-induced root exudate components including 7 amino acids(L-Lysine, L-Methionine, L-Glutamine, L-Leucine, L-Norleucine, L-Isoleucine, and L-Valine), 1 organic acid(6-Aminocaproic acid) and 1 sugar(D-Melezitose) could significantly induce the chemotactic response of A21-4.

**Figure. 4.**
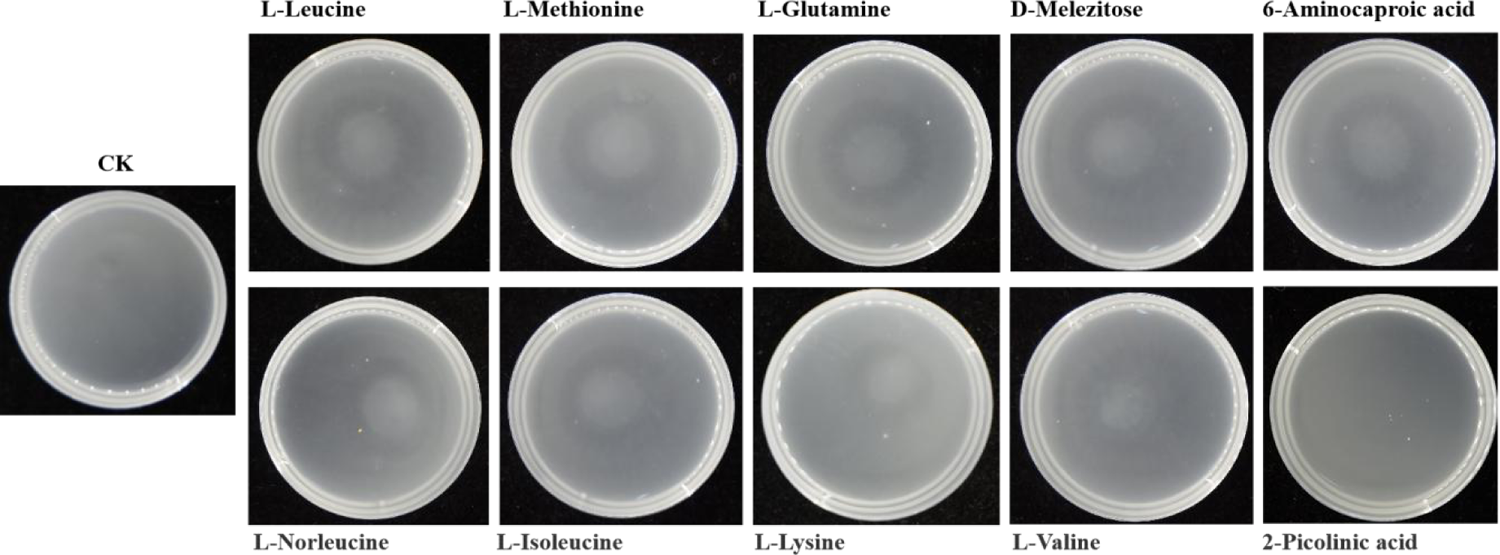
Effect of root exudate components on the qualitative chemotactic response of A21-4. 100 mM of each root exudate component was added to the center of the plate containing viscous cell suspension of A21-4. An equal volume of chemotaxis buffer was used as the control. The plates were placed gently at room temperature for static culture to assess whether a ring formed in the plate.

**Figure. 5.**
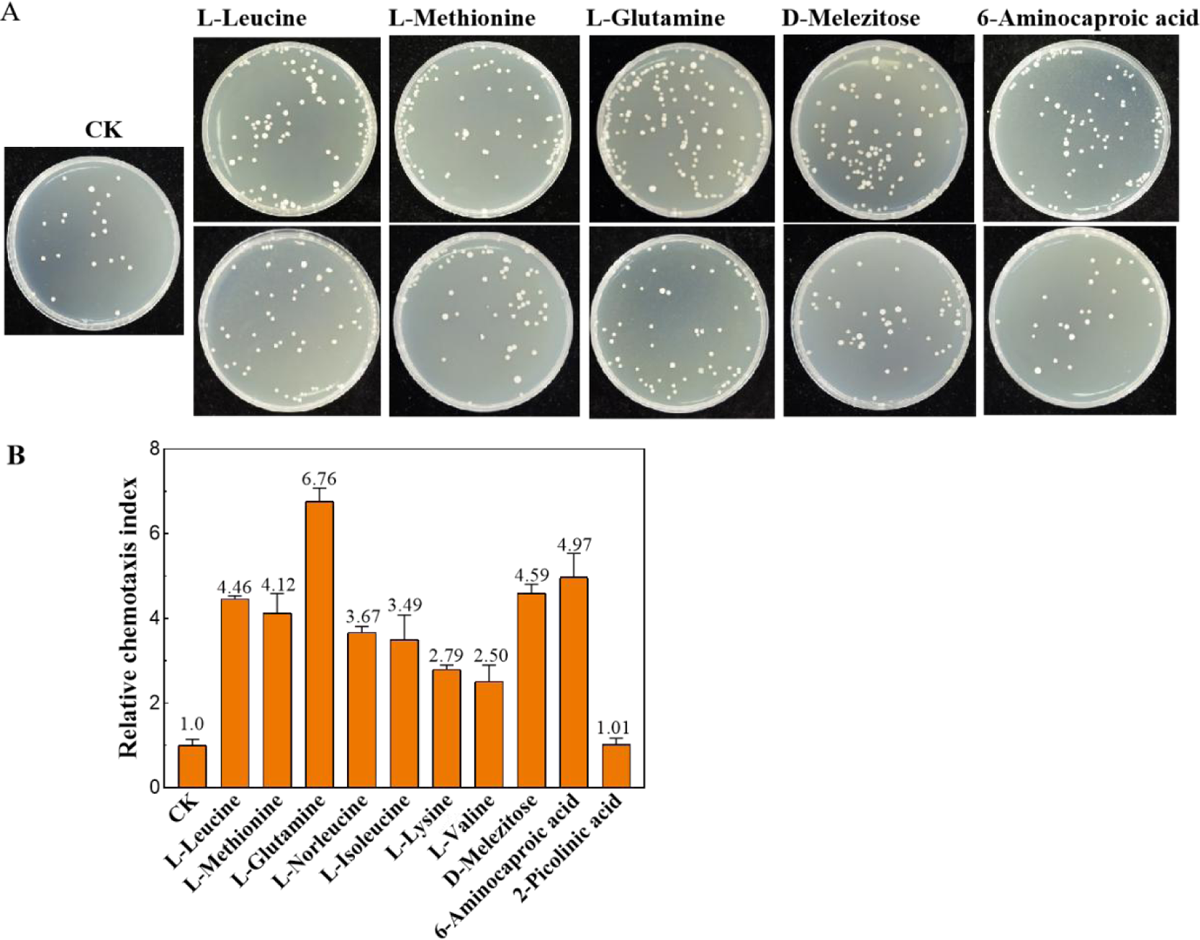
Effect of root exudate components on the quantitative chemotactic response of A21-4. A. Representative image of Colony Forming Units on of A21-4 on TSA medium plates. The bacteria number in the suspensions was obtained by a serial dilution plate and counted after incubation at 28 °C for 48 h. B. Chemotaxis of A21-4 toward different root exudate components. Relative chemotaxis index is calculated based on the ratio of the colony-forming units in response to the different root exudate components to the of the control (sterile water). An RCI ratio ≥ 2 is considered significant.

### 3.4 Selected tomato root exudate components affect biofilm formation of A21-4

To evaluate the biofilm formation of *S. plymuthica* A21-4 in response to selected tomato root exudate components, the methods of crystal violet staining and OD value determination were performed to quantitatively and qualitatively analyze the biofilm formation capacity of A21-4 in different treatments. The results showed that except for 2-Picolinic acid, the rest of 9 selected root exudate components could significantly stimulate the biofilm formation of A21-4. The biofilm biomass of A21-4 in the presence of L-Glutamine, 6-Aminocaproic acid, D-Melezitose, L-Leucine, L-Methionine, L-Norleucine, L-Isoleucine, L-Lysine, and L-Valine was 5. 21, 5. 64, 3. 21, 3. 16, 3. 38, 3. 05, 3. 02, 2. 87, and 2. 73-fold higher than that in the control, respectively. Nevertheless, 2-Picolinic acid also did not markedly induce the biofilm formation of A21-4 compared to the control (Fig. 6). These results indicated that H-R/FR-induced root exudate components including 7 amino acids (L-Glutamine, L-Leucine, L-Isoleucine, L-Lysine, L-Methionine, L-Norleucine and L-Valine), 1 organic acid (6-Aminocaproic acid) and 1 sugar(D-Melezitose) could significantly induce the biofilm formation of A21-4.

**Figure. 6.**
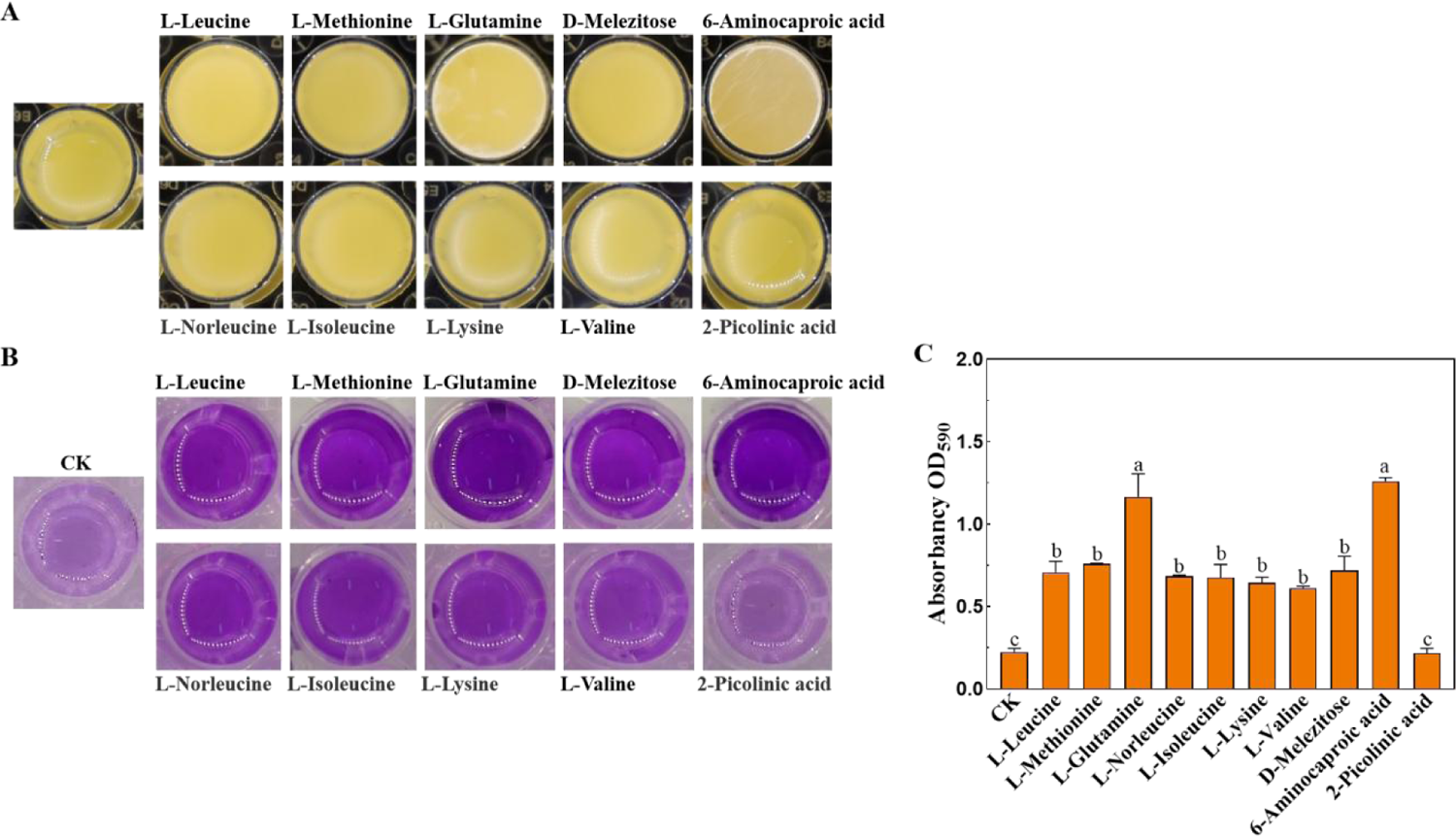
Effect of root exudate components on the biofilm formation of A21-4. A. Microtiter plate assay of biofilm formation of A21-4 in TSB medium containing root exudate components B. Biofilm formation were determined by crystal violetin in the 48-well microtiter plate induced by different components from tomato root exudates. C. OD590 of solubilized crystal violet from the microtiter plate assay. Values are the means ±SD (n=3). The different letters indicate significant difference in the means (*P* < 0. 05, Tukey’s test).

### 3.5 Selected tomato root exudate components enhance the root colonizing and growth promoting capability of A21-4

To find out whether reduced A21-4 root colonization and growth promoting capability in low R/FR light-treated plants were owing to downregulated production of selected tomato root exudate components which further led to the inhibition of chemotaxis and biofilm formation, we hypothesized that A21-4 colonization and growth promoting effect would be rescued by exogenous pretreatment of selected tomato root exudate components. To test these hypotheses we treated both low and high R/FR light-treated plants roots with 100 mM each of reselected tomato root exudate components(L-Leucine, L-Methionine, L-Glutamine, 6-Aminocaproic acid and D-melezitose) for 12 h before inoculating A21-4. When 100 mM L-Leucine, L-Methionine, L-Glutamine, 6-Aminocaproic acid or D-melezitose was pretreated to low R/FR light-treated plants with A21-4 inoculation, plant height and fresh weight were markedly increased over those of the control and A21-4 plants. Moreover, the addition of 100 mM L-Leucine, L-Methionine, L-Glutamine, 6-Aminocaproic acid or D-melezitose further obviously enhanced the plant height and fresh weight of high R/FR ratio-treated plants with A21-4 inoculation when compared with those of plants only with A21-4 inoculation (Fig. 7). Consistent with growth-promoting effects, indeed, each of reselected tomato root exudate components population significantly increased the population density of A21-4 in roots of both high and low R/FR light-treated plants compared with A21-4 treatment, which confirming the roles of reselected tomato root exudate components stimulating colonization of A21-4 (Fig. 8). Collectively, our results strongly demonstrate that high R/FR light-induced root exudate components including L-Leucine, L-Methionine, L-Glutamine, 6-Aminocaproic acid and D-melezitose are essential for growth enhancement and the root colonization of A21-4 in tomato seedlings.

**Figure. 7.**
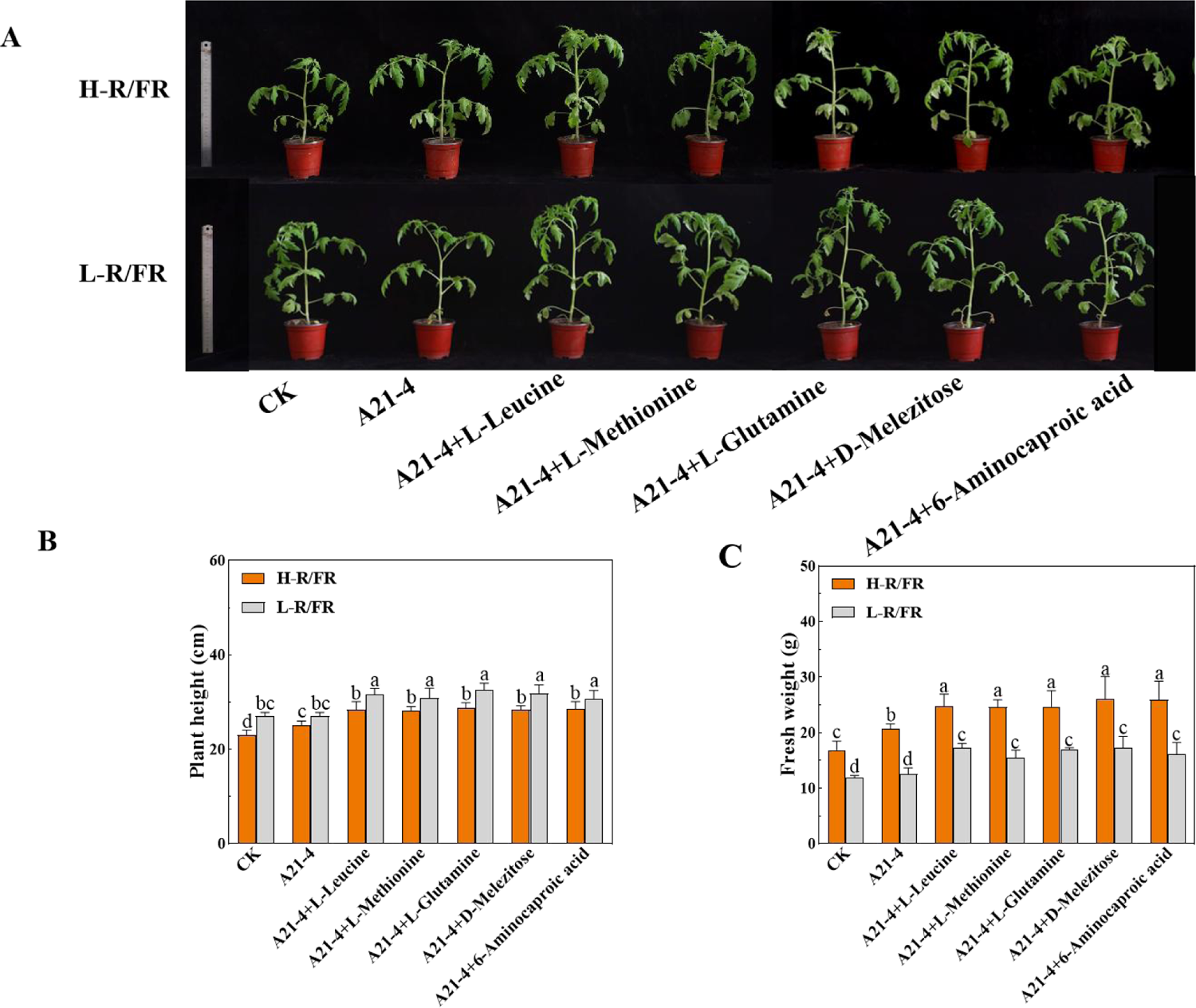
Impact of root exudate components on the growth-promoting effect of A21-4 in tomato seedlings under different R/FR conditions A-C. Plant phenotype, plant height, fresh weight of exogenous root exudate components treated-the tomato plants grown under H-R/FR (2. 5:1) or L-R/FR (1:2. 5) conditions at 7 dpi with (A21-4) or without (CK) inoculation. Values are the means ±SD (n=3). The different letters indicate significant difference in the means (*P* < 0. 05, Tukey’s test).

**Figure. 8.**
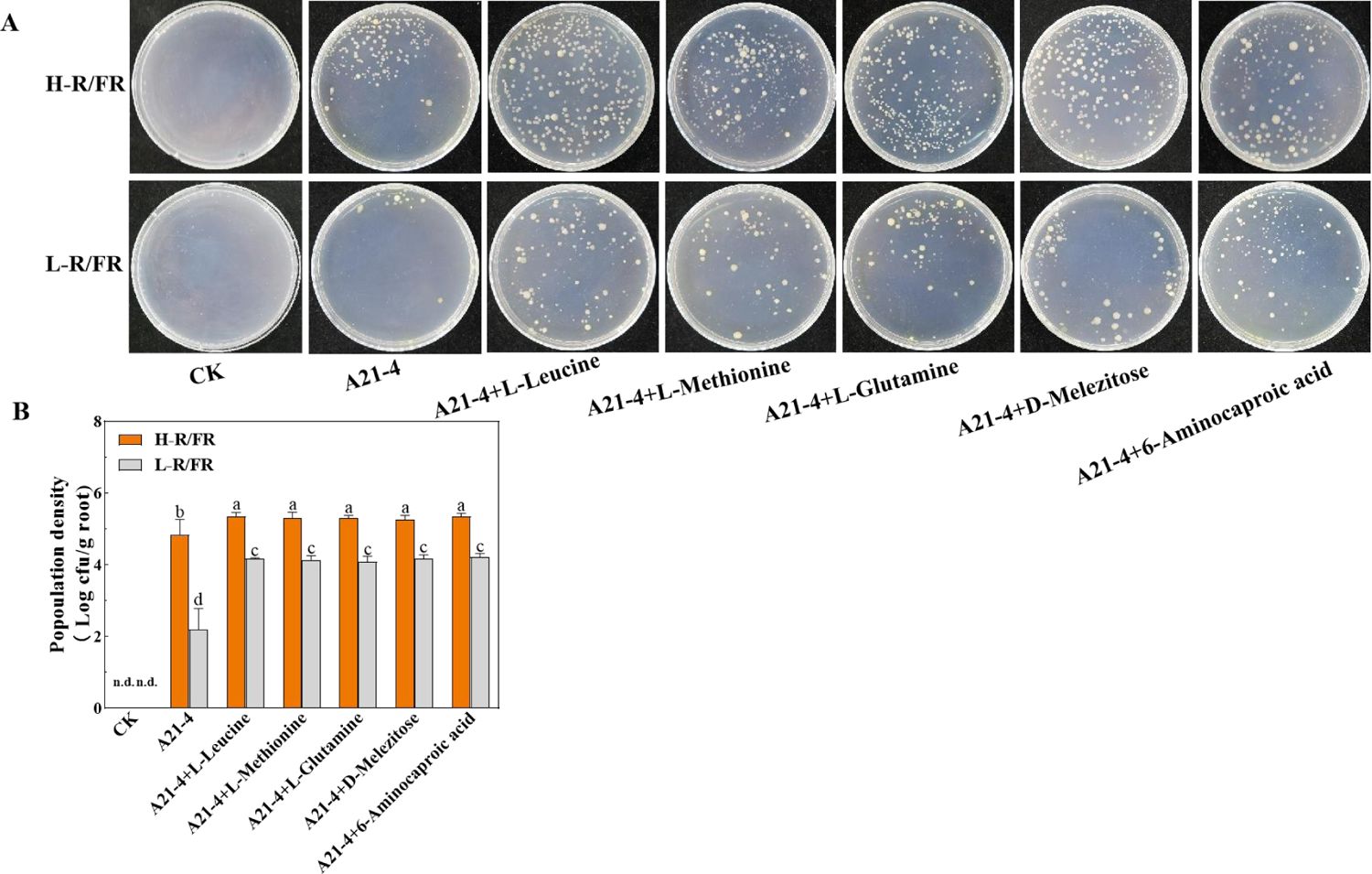
Effect of root exudate components on root colonization of A21-4 in tomato seedlings under different R/FR conditions A. Representative image of Colony Forming Units on of A21-4 on TSA medium plates. B. The population density of the A21-4 on tomato root surfaces was determined by conventional plating method. Values are the means ±SD (n=3). The different letters indicate significant difference in the means (*P* < 0. 05, Tukey’s test).

## 4. Discussion

Successful and stable root colonization of PGPR is crucial for their beneficial functions, whereas the roles and mechanisms of light quality on the interaction of plant and PGPR has not yet been investigated Here, for the first time, we found that high R/FR ratio exposure has been shown to induce the secretion of certain tomato root exudates such as L-Leucine, L-Methionine, L-Glutamine, 6-Aminocaproic acid and D-melezitose by roots, which led to the promotion of the chemotaxis response, biofilm formation and root colonization of A21-4 in tomato seedlings.

PGPR can colonize plant root surfaces to facilitate plant growth and health(Khatoon et al, 2020). In this study, A21-4 promoted tomato seedlings growth, substantially increasing plant height and fresh weight under H-R/FR light but not L-R/FR light condition (Fig. 1A-C). This result offers new evidence for the growth-promoting effect of A21-4 on tomato. Furthermore, our previous studies showed that the strain A21-4 readily colonized on roots of barley, rice, Chinese cabbage, cucumber, watermelon, tomato, and pepper when inoculated on root and seed. Pepper plants with strain A21-4 inoculation had higher seed germination rate, the better growth and greater fruits yield compared with control(Shen et al, 2005). Furthermore, A21-4 also controlled successfully the *P. capsici* induced-Phytophthora blight of pepper(Shen et al, 2007). Therefore, A21-4 has the potential to be developed as a new and important biocontrol agent to promote crop growth and control plant disease.

Light is a key environmental factor that mediates the interaction plant roots and rhizosphere microorganisms(Nagata et al, 2015; Suzuki et al, 2011). For example, the level of AM root colonization in high R/FR light-treated *Lotus japonicus* and tomato was significantly higher than that measured for low R/FR light-treated seedlings(Nagata et al, 2015). Similarly, infection thread formation as well as root nodule formation were significantly inhibited in WT Miyakojima MG20 plants treated by low R/FR light and *phyB* mutants grown under white light(Suzuki et al, 2011). In accordance with these findings, we found that the extent of A21-4 colonization of tomato seedlings grown in high R/FR ratio obviously increased when compared with seedlings grown under low R/FR ratio condition (Fig. 1D). Interestingly, it was recently found that *Bacillus* sp. LC390B-stimulated the growth-promoting effect was disrupted in an *Arabidopsis thaliana phyA/phyB* double mutant, indicating that light perception through PHYA and PHYB is important for plant growth promotion by a beneficial bacterium(Garcia-Cardenas et al, 2023). However, to our knowledge, how light quality modulates root colonization of PGPR such as A21-4 remains unclear.

Root exudates are critical for the root colonization of PGPR(Yuan et al, 2015; Zhang et al, 2014). Plants invest up to 10–40% of their photosynthetically fixed carbon into the rhizosphere as root exudates(Korenblum et al, 2020; Vives-Peris et al, 2020). High-R/FR light preillumination of tomato shoot apex could induce systemic induction of photosynthesis, which was mediated sequentially by phyB, auxin and hydrogen peroxide(Guo et al, 2016). In addition, the low-R/FR light decreased Fe accumulation and photosynthetic electron transport rates in tomato seedlings(Guo et al, 2021). Moreover, low R/FR ratio increased resistance to CO2 diffusion and reduced photosynthetic efficiency in low light grown tomato plants under low light intensities (175μmol m^-2^s^-1^)(Wassenaar et al, 2022). Here, pot experiments that were used to examine the impacts of light quality on the root colonization and growth enhancement of A21-4 in tomato seedlings were also performed at a low light intensities of 140 μmol m^-2^s^-1^, a relatively low light intensity. Thus, we speculated that the R/FR ratio may affected root exudates by partially regulating photosynthesis, thereby affecting the root colonization of A21-4.

Non-targeted metabolite profiling of primary metabolites by UPLC-MS/MS is a powerful technique to link the natural metabolite variation to related plant physiological processes(Danek et al, 2021). Nevertheless, many metabolomics studies have been done on the aboveground parts of the plant, the primary metabolism in root exudates are not deeply investigated(Moenchgesang et al, 2016). Our untargeted metabolic analysis confirmed that the metabolome of root exudates of A21-4-inoculated tomato plants grown under H-R/FR is significantly different from that of A21-4-inoculated roots grown under L-R/FR. Sugars, amino acids, organic acids are considered to be the main attractants to induce chemotaxis of PGPR(Moenchgesang et al, 2016; Singh and Arora, 2001). Consequently, we focused on the three kinds of primary root exudates which were up-regulated by high R/FR light compared with low R/FR light. PCA and OPLS-DA indicated that R/FR ratio exerted an important influence on the composition and quantity of root exudates (Fig. 2). The present study demonstrated that the contents of 1 sugar, 21 organic acids and 31 amino acids in root exudates of tomato seedlings with A21-4 inoculation were significantly higher under high R/FR light than those under low R/FR light (Fig. 3).

Chemotaxis and biofilm formation are recognized as two pivotal and continuous steps for PGPR to effectively colonize on plant roots and exert the beneficial roles(Huang et al, 2022). In view of the availability of different commercially root exudate components and repeatability between different samples, we selected 7 amino acids(L-Glutamine, L-Isoleucine, L-Lysine, L-Leucine, L-Methionine, L-Norleucine and L-Valine), 2 organic acids(6-Aminocaproic acid and 2-Picolinic acid) and 1 sugar(D-Melezitose) to explore their role in chemotaxis and biofilm formation of A21-4. As expected, except for 2-Picolinic acid, the remaining nine selected root exudate components had a good effect to activate the chemotaxis and biofilm formation of A21-4. Chemotaxis effects of selected root exudate components on A21-4 followed the lists of L-Glutamine > 6-Aminocaproic acid > D-Melezitose >L-Leucine>L-Methionine>L-Norleucine>L-Isoleucine>L-Lysine>L-Valine (Fig. 4 and 5). While the ability of selected root exudate components to promote biofilm formation in descending order was: 6-Aminocaproic acid > L-Glutamine > L-Methionine> D-Melezitose > Leucine> L-Norleucine>L-Isoleucine>L-Lysine>L-Valine (Fig. 6). Interestingly, our results are consistent with previous findings that L-Isoleucine, L-Methionine, L-Norleucine, L-Lysine and L-Valine were also shown to be attractive to *Rhizobium lupini* H13-3(Yen et al, 2012). Conversely, L-Methionine was reported to inhibit the chemotaxis of *Bacillus subtilis*, like *Escherichia coli* and *Salmonella typhimurium*(Ordal, 1976). Here, 6-Aminocaproic acid, L-Glutamine and D-melezitose were significantly better attractants of A21-4, which had not been described in other PGPR before. Both chemoattractants and chemoeffectors contribute to the overall chemotaxis to root exudates. For example, CtaA, CtaB, and CtaC are major chemotaxis sensory proteins of the strain *P. fluorescens* Pf0-1 and *ctaA ctaB ctaC* mutant failed in chemotaxis to amino acids exhibited a noticeably decreased ability to colonize tomato root(Oku et al, 2012). In the present study, chemotaxis sensory proteins of A21-4 for selected root exudate components and their genes were not designated. Thus, it will be interesting to study the key chemoreceptors sensing dominant attractants and mediating recruitment of A21-4 in the future.

Promoting the root colonization process by PGPR is essential for the use of these strains in agriculture. In this study, low R/FR light condition was suboptimal for A21-4 colonization and growth enhancement, while the rescue of A21-4 colonization by exogenous pretreatment of tomato root exudate components in low R/FR light-treated plants were very effective (Fig. 7 and 8). Thus, on one hand, high R/FR light ratios during the day or nighttime supplementation of R light can be used to improve A21-4 colonization under greenhouse conditions; on the other hand, exogenous pretreatment of tomato root exudate components prior to A21-4 inoculation can be used to enhance A21-4 colonization in the field. In addition, we should also pay attention to control the crop planting density in field and greenhouse, in order to avoid that seedlings experience low R/FR ratio when grown at high planting density, which further lead to ineffective rhizosphere colonization of A21-4.

## 5. Conclusion

In summary, high R/FR light promotes and low R/FR light inhibited the root colonization and growth-promoting effect of A21-4 on tomato. High R/FR light treated tomato seedlings secreted L-Glutamine, L-Leucine, L-Methionine, 6-Aminocaproic acid and D-Melezitose, which stimulated the chemotaxis response and biofilm formation of A21-4. Artificial root exudate supplementation can significantly improve the strain A21-4 to colonize the surface of tomato roots. These findings lay a foundation for the effective combined application of light-emitting diodes and PGPR, thereby availably promoting plant growth and health and minimizing the use of chemical fertilizers, pesticides in green agricultural production.

## Authors contributions

Zhixin Guo and Shunshan Shen designed the research and finished the manuscript. Zhixin Guo, Yanping Qin, Jingli Lv, Xiaojie Wang, Ting Ye, Xiaoxing Dong, Nanshan Du and Tao Zhang conducted the experiments. Zhixin Guo, Jingli Lv and Fengzhi Piao analyzed the data. Han Dong, Yanping Qin and Shunshan Shen revised the article. All authors read and approved the final version of the manuscript.

## Conflicts of interest

The authors declare no conflict of interest.

## Acknowledgements

This work was supported by National Natural Science Foundation of China(NSFC)(32002118; 32272794; 32202579), China Agriculture Research System (CARS-23-B03), and Key scientific research project of Henan Province(22A210008).

## Data Availability Statement

All data generated from the study appear in the submitted article.

## Notes

### Competing Interest Statement

The authors have declared no competing interest.

